# Tools and Resources: Remote, automated and MRI-compatible administration of interoceptive inspiratory resistive loading

**DOI:** 10.1101/2019.12.09.869511

**Authors:** Sebastian W Rieger, Klaas Enno Stephan, Olivia K. Faull

## Abstract

Research on how humans perceive sensory inputs from their bodies (‘interoception’) has been rapidly gaining momentum, with interest across a host of disciplines from physiology through to psychiatry. However, studying interoceptive processes is not without significant challenges, and many methods utilised to access internal states have been largely devoted to capturing and relating naturally-occurring variations in interoceptive signals (such as heartbeats) to measures of how the brain processes these signals. An alternative procedure involves the controlled perturbation of specific interoceptive axes. This is challenging because it requires non-invasive interventions that can be repeated many times within a subject and that are potent but safe. Here we present an effective methodology for instigating these perturbations within the breathing domain. We describe a custom-built circuitry that is capable of delivering inspiratory resistive loads automatically and precisely. Importantly, our approach is compatible with magnetic resonance imaging environments, allowing for the administration of complicated experimental designs in neuroimaging as increasingly required within developing fields such as computational psychiatry/psychosomatics. We describe the experimental setup for both the control and monitoring of the inspiratory resistive loads, and demonstrate its possible utilities within different study designs. This methodology represents an important step forward from the previously utilised, manually-controlled resistive loading setups, which present significant experimental burdens with prolonged and/or complicated sequences of breathing stimuli.

## Introduction

A fundamental aspect of complex beings is the sensation, perception and control of the physical body. ‘Interoception’ can be considered as the processing of sensory inputs from the body by the nervous system across both subconscious and conscious domains^1^. However, surprisingly little is known about the dynamic interplay between the brain and body beyond homeostatic reflex control^2-4^, and the burgeoning field of interoceptive research has its sights set on understanding brain-body axes for potential applicability to emotions, decision making, consciousness and mental health (recently reviewed by Khalsa and colleagues^1^), to name a few. In this article we propose a significant methodological advancement regarding the experimental requirements for investigating interoception in the respiratory domain.

In order to study the brain-body interface across interoception, we need to equip ourselves with tools for capturing (and potentially inducing) perturbations in interoceptive experiences. This may take the form of measuring and modelling natural fluctuations that occur within and across interoceptive domains, such as variations in heart rate^5,6^, breath-to-breath variability in respiration^7,8^, or irregularities across the gastric cycle^9^. Alternatively, we can consider actively invoking changes in these systems in a controlled, timely and reversible manner. Such active disturbances are critical for testing hypotheses about the principles that govern interoception. For example, current theories postulate that interoception obeys the same hierarchical Bayesian principles as exteroception ^2-4,10^. A critical prediction of these theories concerns the occurrence of specific prediction error signals (interoceptive surprise), e.g. reflected by activity of the insula, in response to unexpected changes in sensory inputs from the body. Testing this prediction requires experimental perturbations that elicit controlled prediction errors in a trial-by-trial fashion. Active perturbations of bodily states are also important for clinical applications^11,12^, not least in computational psychiatry/psychosomatics where model-based assessment of interoceptive surprise in individual patients plays a central role in proposals for differential diagnostics^3,4^.

Methods to induce controlled perturbations of interoception have so far been trialled in the cardiac domain (via intra-venous administration of isoproterenol^13^), with breathing (via inspiratory and/or expiratory resistance^12,14-17^), with gastric and bladder disturbances (via inflation of inserted balloons^18^, or via baroreceptor manipulation^19^). While pharmacologically-induced cardiac alterations induce strong effects and appeal to the wide body of research on natural fluctuations in interoceptive heart-related signals^20-25^, the time-related decays required for returning to baseline as well as the burden and risks associated with repeated pharmacological interventions within the same subject pose restrictions on widespread application of this method. Similarly, the use of both gastric/bladder balloons and baroreceptor manipulations face challenges with regard to patient comfort and acceptance.

Here, we pursue a different approach for perturbing interoception in a fashion that is controlled, safe, repeatable, and not too inconvenient for human participants. We focus on respiration, presenting an advanced circuitry for the automated administration of inspiratory resistances. This circuitry builds on a previously-published magnetic resonance imaging (MRI)-compatible inspiratory resistance circuit^16,26,27^, incorporating computer-controlled solenoid valves for timely commencement and elimination of resistance, and an electronically-controlled, flow-mediated inspiratory valve device (POWERbreathe, IMT Technologies Ltd., Birmingham, UK) to allow for adjustable maximal resistances. While the previous methodology has been employed to measure the brain activity (via functional magnetic resonance imaging, or fMRI) related to the conditioned anticipation and perception of inspiratory resistive loads, these advances significantly aid the development of complicated protocols required for a more detailed study of dynamic brain-body interactions^28^. These protocol requirements often call for accurately controlled, repeatable and changeable administration of interoceptive perturbations, which are essential to amass the necessary statistical evidence for quantifying subject-specific processes involved in interoception such as dynamic predictions and learning^29^.

## Methods

### Previous inspiratory resistance administration circuitry

The previous circuitry that has been employed to manually administer inspiratory resistances within an MRI scanner^16,26,27^ is outlined in Figure 1A. In this experimental setup, compressed and humidified medical air was delivered to the participant via a breathing system, whereby air flow was maintained at a rate to adequately allow free breathing and access to an available air reservoir (within a reservoir bag) of 2 L. When a (conditioned) visual cue appeared on the screen, the delivery of compressed air was manually halted via closure of the corresponding air flowmeters, allowing the reservoir bag to empty over the course of approximately 3-8 seconds (anticipation period), followed by the application of inspiratory resistance once the air reservoir was empty. Additionally, delivery of supplementary gas mixtures (oxygen (O_2_) and a carbon dioxide (CO_2_) mix of 10% CO_2_, 21% O_2_, balance nitrogen) was supplied to the participant, allowing for periods of manual gas mixture administration (during rest periods) to decorrelate changes in expired CO_2_ and O_2_ from periods of inspiratory resistance.

**Figure 1.**
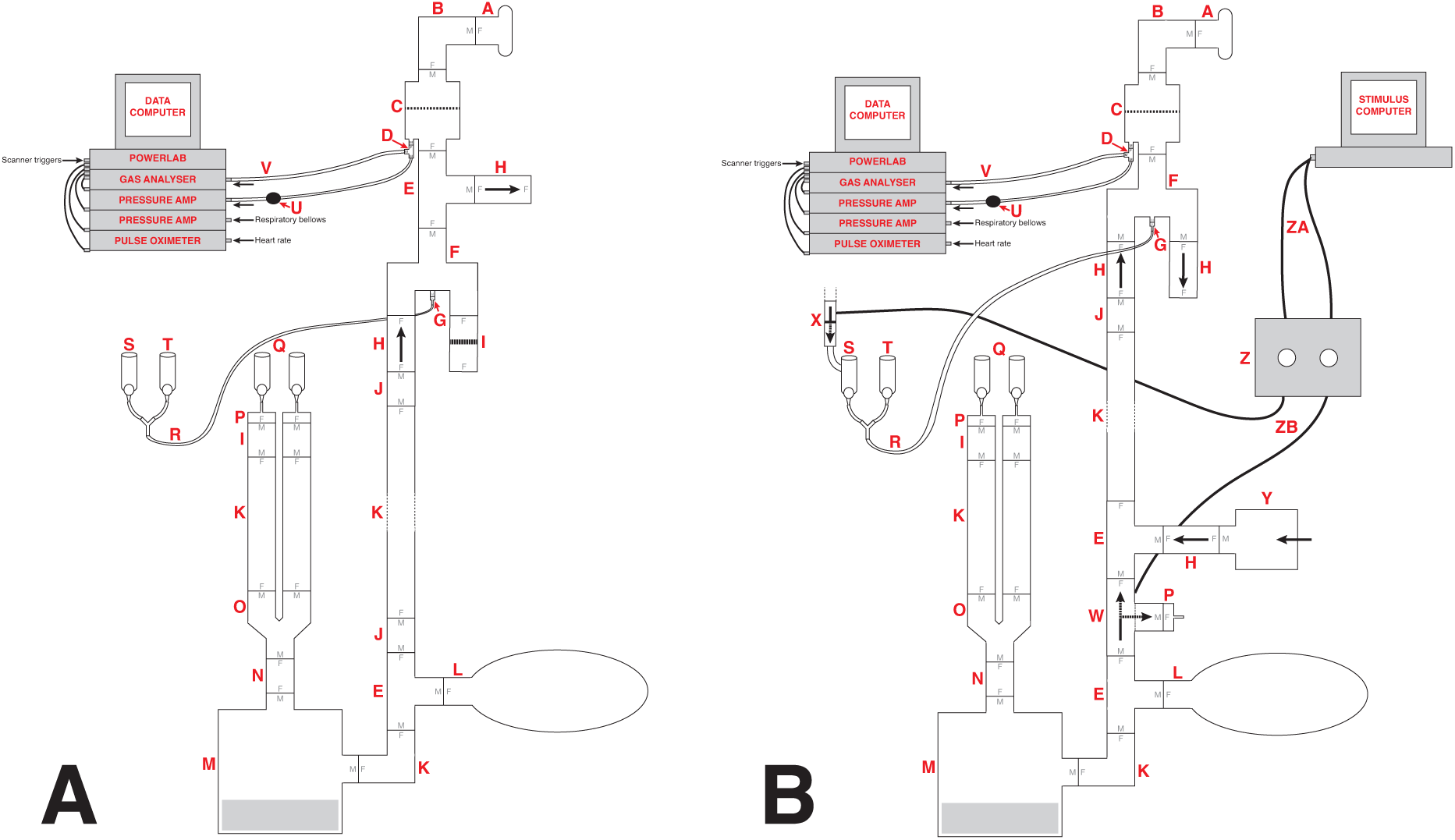
Schematic of the previously-utilised inspiratory resistance circuit^16,26^ (presented in panel A) and the new circuit design (panel B) that allows remote administrations of inspiratory resistance. In both systems, medical air is supplied to the subject, with a reservoir of 2 L. Excess flow and expiration escapes through a one-way valve (labelled H), close to the mouth to minimise rebreathing. A diving mouthpiece (labelled A) is connected to a bacterial and viral filter (labelled C), and sampling lines connect to a pressure transducer (labelled U) and amplifier (Pressure transducer indicator, PK Morgan Ltd, Kent, UK) for inspiratory pressure readings, and to a gas analyser (via sampling line labelled V) (Gas Analyser; ADInstruments Ltd, Oxford, United Kingdom) for respiratory gases. In diagram A, resistive inspiratory loading is induced by discontinuing the delivery of medical air (via the flowmeter and emptying of the reservoir bag), forcing the subject to draw air through the resistor (porous glass disc labelled I). In diagram B, resistive inspiratory loading is automatically achieved via the stimulus computer, whereby signals are sent through the parallel port to control valve 1 (labelled W) to redirect the supply of medical air to vent to the environment, forcing the subject to draw air through the POWERbreathe device (labelled Y). Periodically throughout scanning, small boluses of additional carbon dioxide (CO_2_) can be administered through manual control of the CO_2_ flowmeter (labelled S) in diagram A, or automatic control via valve 2 (labelled X) in diagram B, to raise the partial pressure of end-tidal CO_2_ (P_ET_CO_2_) to match the P_ET_CO_2_ rise induced by inspiratory loading periods. A final flowmeter (labelled T) is available for manual input of additional oxygen (O_2_) to the system. A full list of the labelled component parts can be found in the supplementary material.

### Solenoid valves

Within the new inspiratory resistance circuitry design (Figure 1B), two solenoid valves have been placed to allow for the automated and immediate application of inspiratory resistances, and administer small boluses of CO_2_ in a predetermined manner. Valve 1 (W in Figure 1B) is a wide-bore, direct-acting, 3/2-way solenoid valve (3/918-24/1002/R370-GN N.O. 24V DC, Buschjost Magnetventile GmbH & Co. KG, Germany) fitted with brass stem adaptors (MM052206N, John Guest Ltd.) to match the diameter of the air supply tubing, reducing the influence of any added baseline resistance to breathing. This valve is of the “normally open” type and will in its default state allow air to flow from the supply towards the mouthpiece. When actuated, the valve shuts off the port which supplies the mouthpiece and so initiates inspiratory loading. At the same time, the valve vents the medical air supply to the atmosphere to prevent overpressure. Valve 2 (X in Figure 1B) is a normally closed direct-acting 2/2-way valve (type 6013, Bürkert GmbH & Co. KG, Germany) that is inserted into the gas supply line for the CO_2_. This valve is closed by default, and when activated will open to allow the administration of the gas mixture for any adjustable length of time.

### Solenoid valve control box

The solenoid valves are operated via a custom-built control circuitry box (Figure 2), which can be interfaced with a presentation software program such as PsychToolBox^30^, or activated manually via a button press on the control box. The valve controller uses a 24V DC 5A power supply switched using optically coupled solid state (photoMOS) relays (AQZ102, Panasonic). Each output is connected to the supply via two parallel relays, one of which can be operated via a pushbutton, while the other one is energised directly from the TTL compatible control input. This allows standalone manual operation (without a PC connected) as well as manual override (i.e. manual operation with a PC connected), while ensuring the TTL inputs remain galvanically separated from the power supply and valve circuits. Kickback protection is achieved by the use of suitable solenoid valve leads (40881 Series, Murrelektronik).

**Figure 2.**
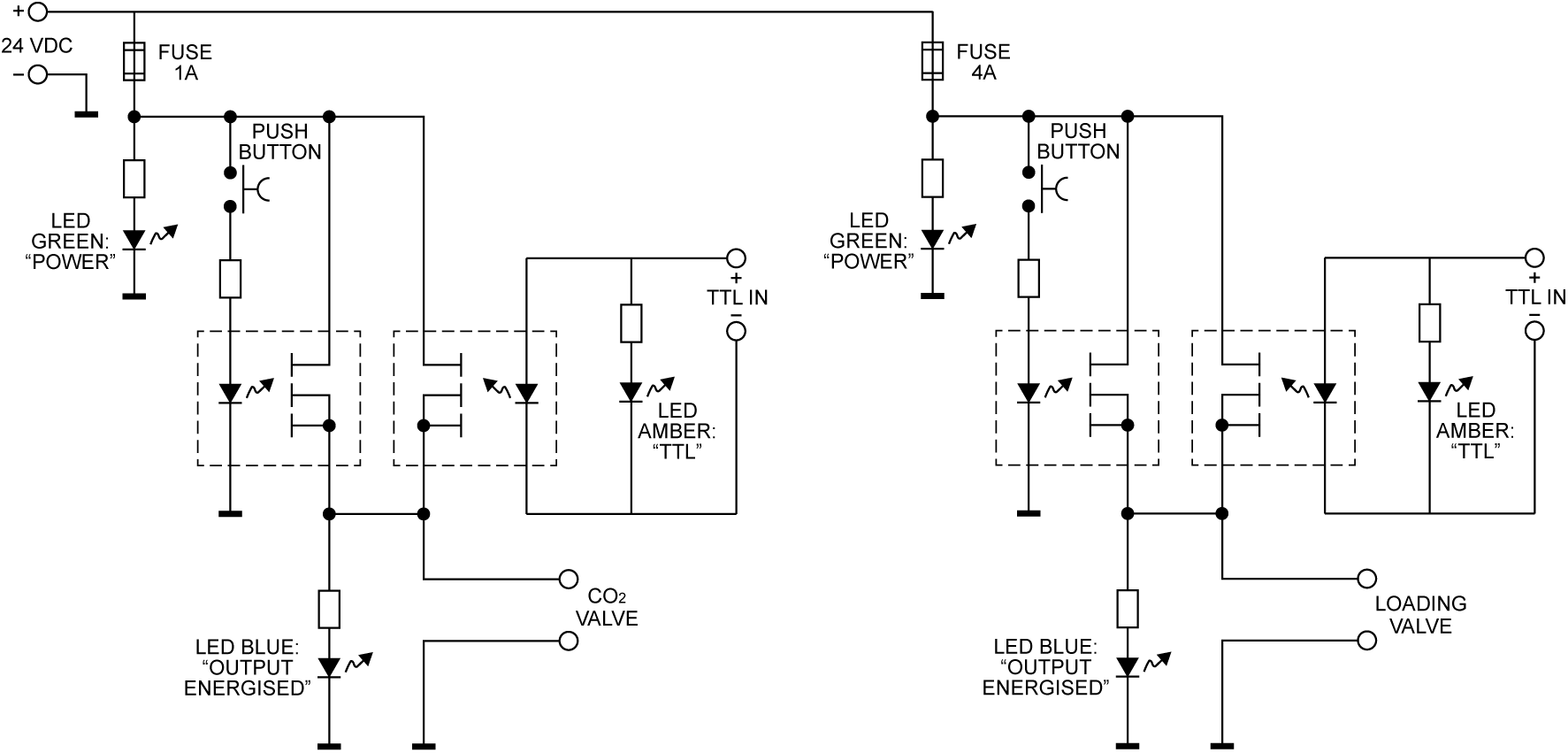
The valve controller circuit. Each of the two solenoid valves is switched using a pair of Panasonic AQZ102 optoFET relays connected in parallel, allowing them to be energised individually, either manually via the pushbutton, or from a TTL signal, while ensuring the TTL input remains galvanically separated from the control circuit. Power is supplied from a standard 24VDC power supply (Meanwell GST90A24-P1M, not shown). For additional protection from transients at the outputs, the valves are connected using cable assemblies with built in suppression (Murrelektronik 7000-40881-6360200, not shown).

To interface with PsychToolBox (or other presentation programs), simple commands can be sent via the parallel port of the presentation computer and be fed directly into the control box. This feature allows for completely automated resistance administration, which could either be using a predetermined, randomised or online-updating sequence of stimuli. Sample code for control via the parallel port is provided in the supplementary material.

### Adjustable maximal resistance

To allow for an adjustable maximal inspiratory resistance, the porous glass disc previously used to evoke an inspiratory resistance (Figure 1A) has been replaced with an electronically-controlled, flow-mediated resistance valve (POWERbreathe, IMT Technologies Ltd., Birmingham, UK). This device can be set to an inspiratory resistance between 3 and 200 cmH_2_O, and when the pressure generated by the participant inspiring against the POWERbreathe valve exceeds this set resistance, the valve is released.

## Results

### Resistance administration

In Figure 3 we demonstrate the physiological traces recorded during the repeated activation of valve 1, and subsequent inspiratory resistance applications for a set value (here we applied a resistance value of 55 cmH_2_O, which was determined by calculating 70% of the participant’s maximal inspiratory pressure of 79 cmH_2_O. The participant was instructed to maintain normal breathing depth and rate during the resistive loading periods, and subsequently generated a pressure of approximately 8 – 10 cmH_2_O in each breath. In Figure 4, we demonstrate the physiological traces recorded during the repeated activation of valve 1 with 5 cmH_2_O graded increases in set pressure from 5 – 30 cmH_2_O.

**Figure 3.**
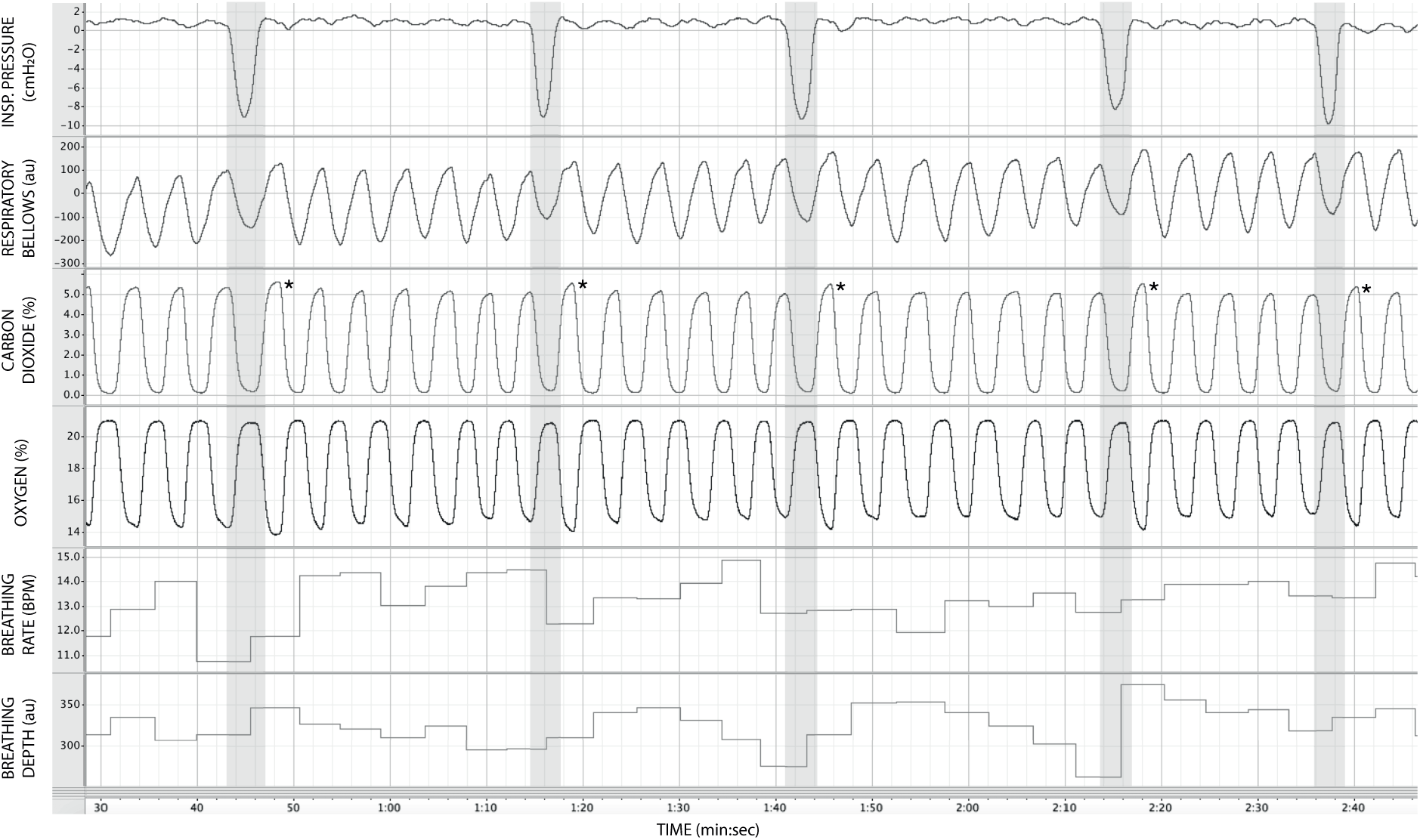
Physiological traces recorded during the application of five inspiratory resistance periods (grey areas), where the maximal inspiratory resistance was set at 55 cmH_2_O. Asterisks (*) denote an increase in expired pressure of end-tidal carbon dioxide (P_ET_CO_2_ – determined by the peak values at the end of each expiration in the CO_2_ trace) immediately following each inspiratory resistance period.

**Figure 4.**
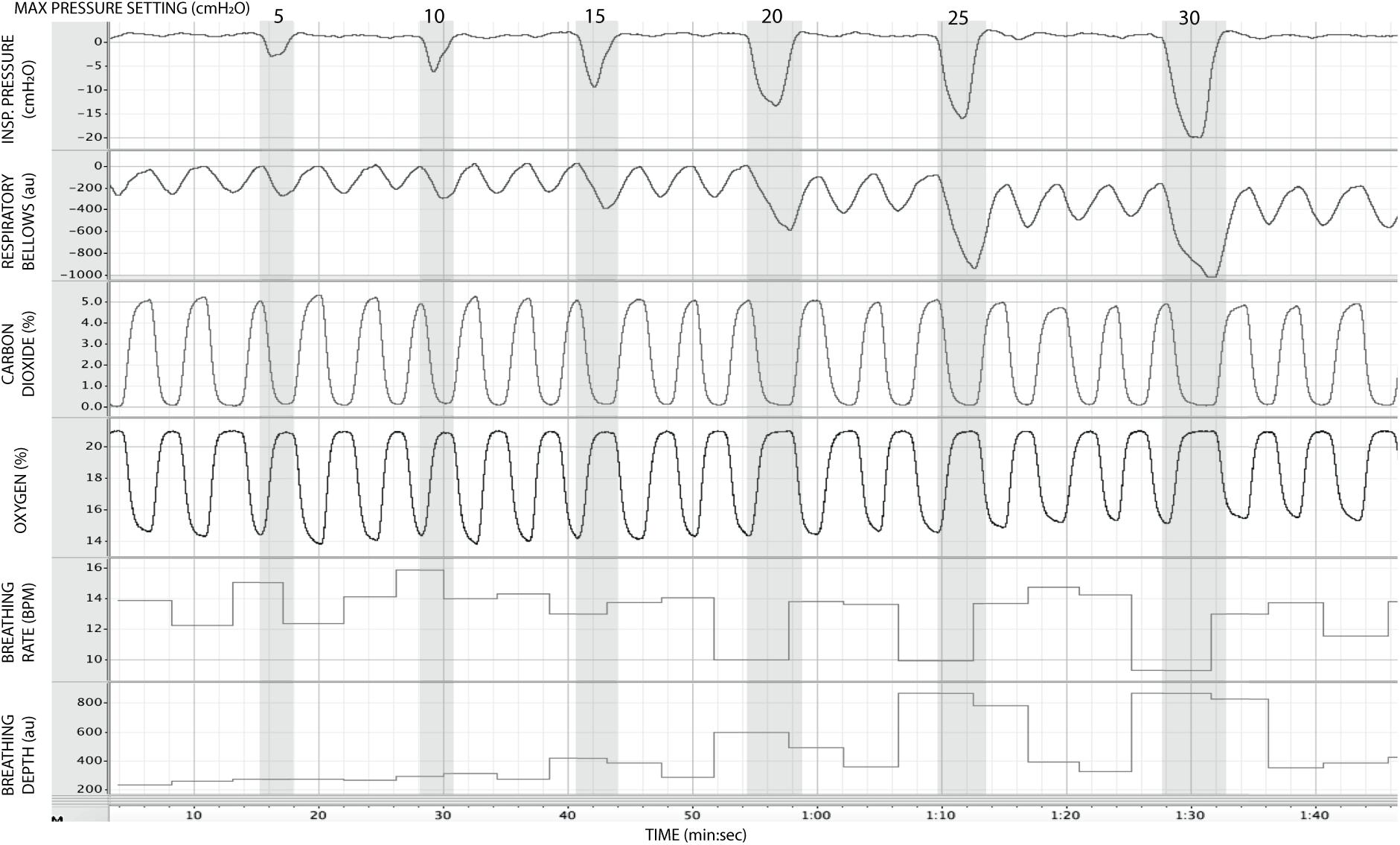
Physiological traces recorded during the application of five inspiratory resistance periods (grey areas), where the level of inspiratory resistance was graded at intervals of 5 cmH_2_O between 5 – 30 cmH_2_O.

### Administration of additional gas mixtures

In Figure 5 we demonstrate the physiological traces recorded during repeated activation of valve 2, and the subsequent administration of small boluses of elevated CO_2_ (achieved here using a short (< 1 second) administration of a mixture containing 20% CO_2_, 21% O_2_ and the balance nitrogen). Importantly, the end-tidal values of expired CO_2_ (the peak value at the end of expiration, highlighted in Figure 5) can be seen to elevate and then recede over the course of 2 – 3 breaths, which could be extended by specifying a longer duration of administration.

**Figure 5.**
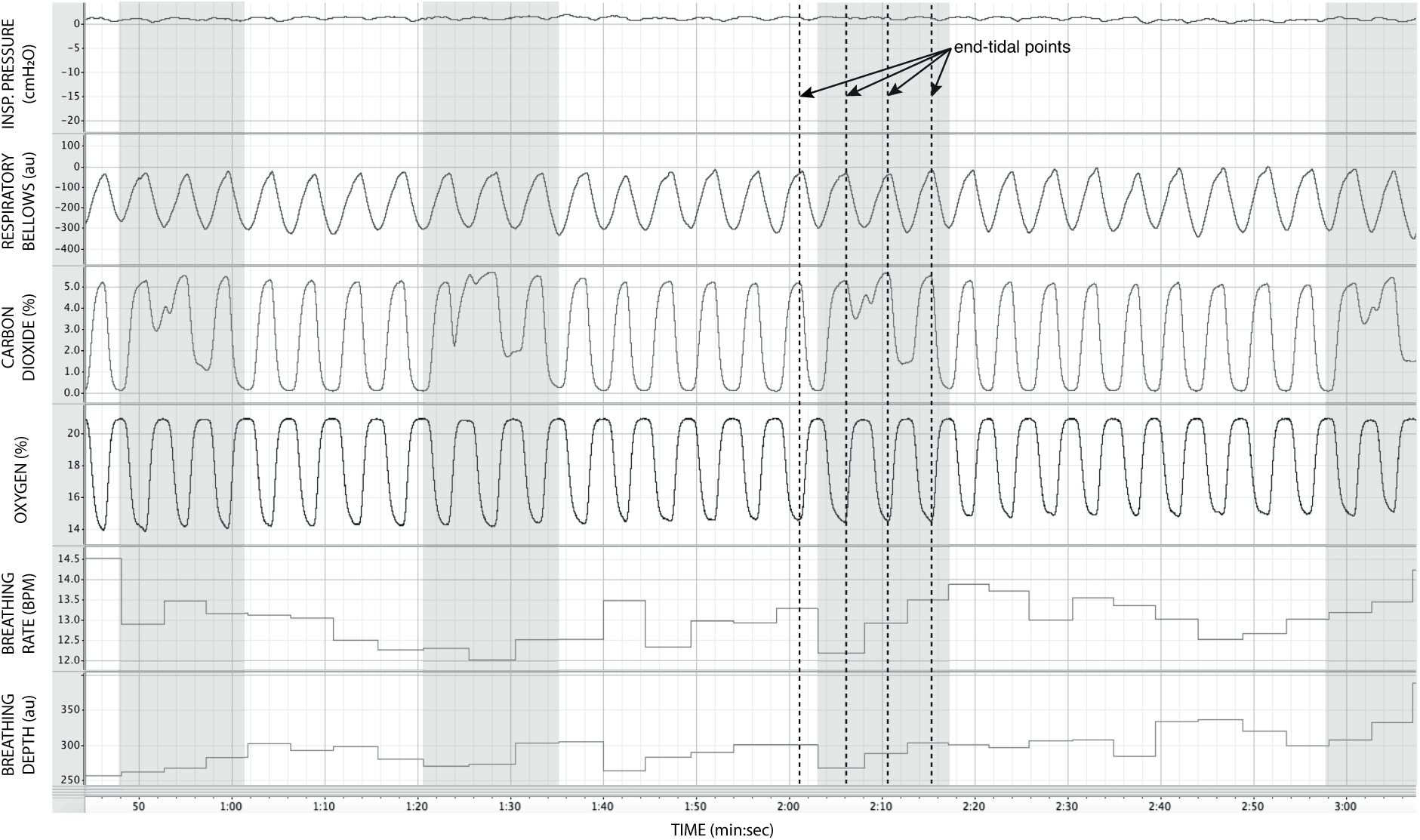
Physiological traces recorded during the application of four boluses of a carbon dioxide mixture (grey areas: 20% CO_2_, 21% O_2_ and the balance nitrogen) added to the medical air supply. Dotted lines represent example end-expiratory points, from which end-tidal CO_2_ and O_2_ values would be taken.

## Discussion

In this report we have presented a custom-designed circuitry to administer automated, remote, timely and repeated inspiratory resistances suitable for an MR environment. In this design we have included the use of solenoid valves to control the administration of both inspiratory resistance and boluses of additional gas mixtures, which can be triggered either from a stimulus computer or by a manual button press on a custom-built circuitry box. This design represents a significant improvement from previously-published methodologies^16,26,27^, whereby inspiratory resistances were administered via manual closing of flowmeters and emptying of reservoir bags, both of which present substantial obstacles as the necessity of more complicated experimental designs beckons. As outlined in the Introduction, methodologies such as the one described in this paper represent an essential step in our progression towards understanding the links between brain and body, and the importance of interoceptive processes across a wide range of neuroscientific domains.

The circuit design presented here has been specifically tailored to cater for the demands of an MRI environment. In this setting, where all equipment within the scanner room is required to be non-magnetic, the solenoid valves, electrical circuitry, gas flow control and inspiratory resistance device can all be kept within the scanner control room, with flexible lengths of tubing allowing for the remote delivery of air supply and gas mixtures to scanners of variable distances from a control room port. All equipment within the scanner room is made of plastic or non-ferromagnetic materials, and both participants and the circuit are protected via a disposable, single-use bacterial and viral filter. Additionally, the sampling of gas and inspiratory pressures is fed back via sampling lines to monitoring devices that are also housed within the control room. To account for any gas sampling time-delays caused by the length of sampling line required, delays can simply be measured and the temporal shift incorporated into the analysis of any recorded physiological traces.

The advances made in this circuit design provide a platform from which a number of inspiratory resistance protocols are compatible. One of the simpler methodologies would be to employ an on/off binary protocol, where the maximal resistance on the POWERbreathe is set to a percentage of each subject’s maximum inspiratory pressure (which can be tested using the POWERbreathe device). This protocol should allow a very clear perceptual distinction between the presence and absence of an inspiratory resistance, and could then be programmed and automated to fulfil the requirements of the research question at hand. Furthermore, as the pressure generated on each trial results from the inspiratory effort produced by the participant, either visual biofeedback could be employed to ensure pressure consistency in each trial, or the natural variability in both inspiratory pressure and perceptual ratings across trials could be used to tease apart differences in physiological vs. perceptual brain activity. Alternatively, graded levels of inspiratory resistance could be employed via a pre-determined protocol input to the POWERbreathe, although visual feedback may be a necessity here to ensure participants reach the desired inspiratory pressure in each trial. This is exemplified in Figure 4, whereby the distention on the respiratory bellows increases with increasing resistance, demonstrating an increase in the inspiratory depth required to overcome the set resistance. Additionally, it should be noted that the pressure generated at the mouth during inspiratory resistance periods falls below the resistance set at the POWERbreathe (Figure 4). This recorded pressure differential will be proportional to factors such as the inspiratory effort generated by the participant, as well as the flow resistance of the tubing between the participant and the device (in turn influenced by the length of tubing required to reach the participant in the particular experimental setup). Therefore, it is imperative that the inspiratory pressure generated is additionally recorded at the mouth for experiments utilising this design, as a metric of inspiratory effort against the resistance within the specific experimental setup.

The design of this circuit also provides a means to tackle one of the key issues when utilising breathing-related tasks in an fMRI protocol: Namely, the disturbances in blood gas concentrations and their subsequent effects on the blood oxygen level dependent (BOLD) signal that is the fundament of most functional imaging sequences. The partial pressure of CO_2_ (PCO_2_) in the blood is a potent vasodilator, and tasks that induce changes in blood gases via altering natural breathing rhythms can thus either wash-out BOLD signal (via increases in PCO_2_) or induce global grey-matter vasoconstriction (via decreases in PCO_2_) that may interact non-linearly with the neutrally-induced BOLD response^31^. Additionally, changes in the partial pressure of oxygen (PO_2_) in the blood also plays a vasodilatory role, however the magnitude of this response is considerably smaller than that induced by PCO_2_^31^. In this circuit design, we are firstly able to remotely measure the changes in expired CO_2_ and O_2_, whereby the end-tidal values are assumed to be indicative of alveolar gas (which closely parallels arterial carbon dioxide levels under normal ventilation-perfusion matching^32^). Secondly, solenoid-valve controlled boluses of additional CO_2_ can also be delivered whenever required (for example in some rest periods of the protocol), which allows the changes in expired gas to be decorrelated from the breathing task^33^. However, it should be noted that maintenance of stable end-tidal gas measurements via end-tidal forcing^34^ has not currently been incorporated into this system, although this could be integrated in place of the binary solenoid CO_2_ valve if necessary.

## Conclusion

Here we have presented a circuitry that has been designed to automatically administer periods of inspiratory resistance in a remote environment (such as an MRI scanner), allowing the potential instigation of more complex paradigms that probe the relationship between brain and body. However, whilst breathing represents a somewhat more easily accessible avenue in which to study interoception, the changes in blood gas pressures that often result from breathing tasks cannot be overlooked. Therefore, in this design we have incorporated methods for both recording and decorrelating fluctuations in the pressure of expired CO_2_ and O_2_, as a representation of fluctuations in arterial gas pressures. Lastly, the use of an electronically controlled inspiratory resistance device allows for specification of a ceiling of maximal inspiratory resistances, although careful use of biofeedback may be necessary to ensure graded levels of pressure are reached by the participant on each trial if required. We hope that this circuit design may aid the more nuanced study of brain-body interactions and interoception in the respiratory domain.

## Acknowledgements

Olivia Faull is a Marie Skłodowska-Curie Postdoctoral Fellow that is supported by the European Union’s Horizon 2020 research and innovation programme under the Grant Agreement No 793580. Klaas Enno Stephan was supported by the René and Susanne Braginsky Foundation and the University of Zurich.

## Supplementary Material

### 1) Full equipment list for Figure 1 (reproduced below)

#### Equipment list

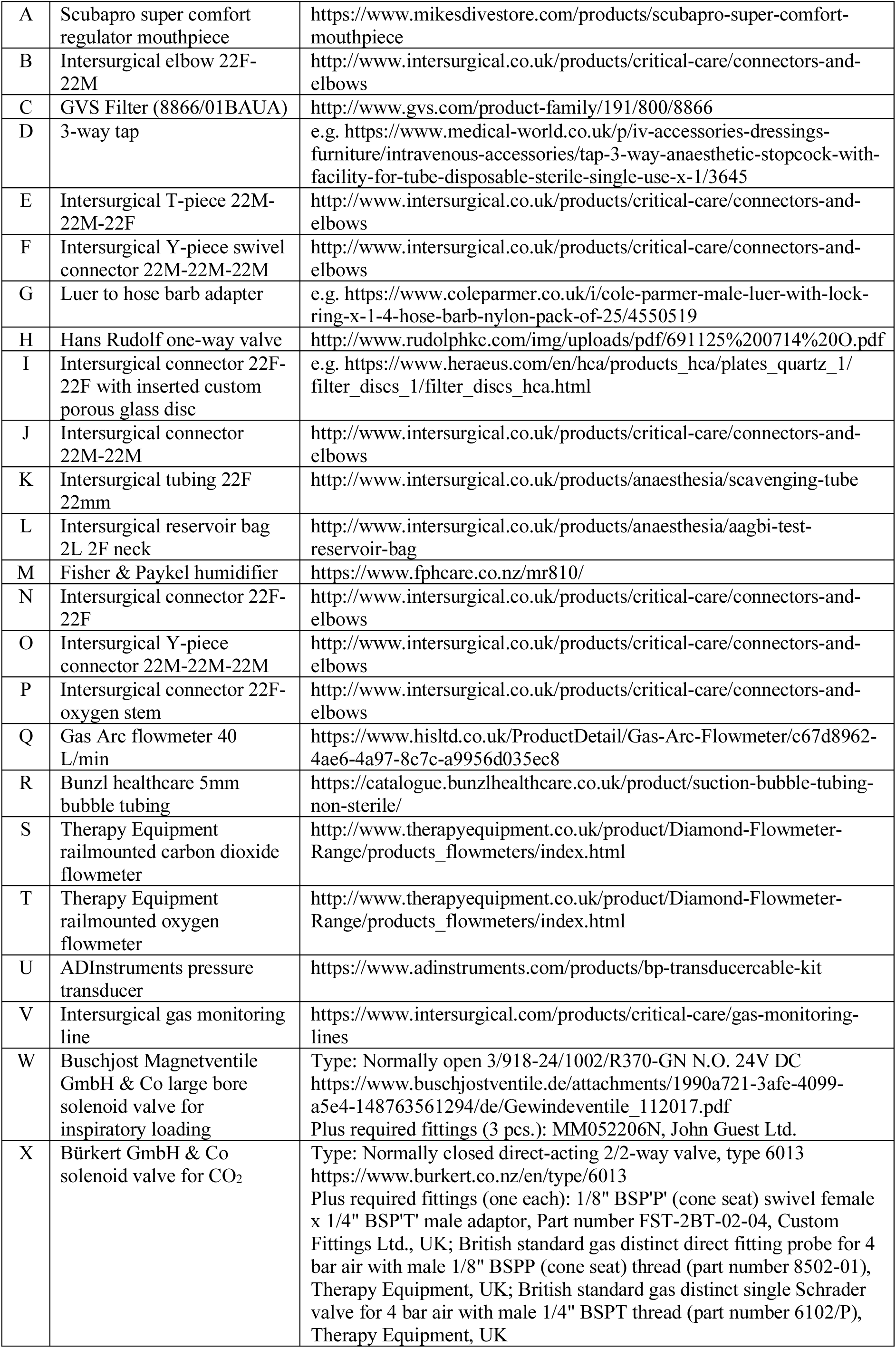
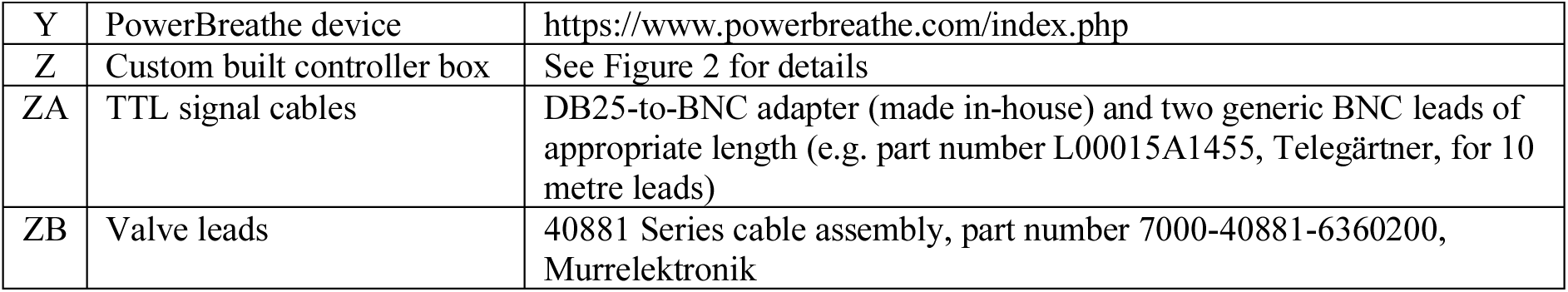

|  |  |  |
| --- | --- | --- |
| A | Scubapro super comfort regulator mouthpiece | https://www.mikesdivestore.com/products/scubapro-super-comfort-mouthpiece |
| B | Intersurgical elbow 22F-22M | http://www.intersurgical.co.uk/products/critical-care/connectors-and-elbows |
| C | GVS Filter (8866/01BAUA) | http://www.gvs.com/product-family/191/800/8866 |
| D | 3-way tap | e.g. https://www.medical-world.co.uk/p/iv-accessories-dressings-furniture/intravenous-accessories/tap-3-way-anaesthetic-stopcock-with-facility-for-tube-disposable-sterile-single-use-x-1/3645 |
| E | Intersurgical T-piece 22M-22M-22F | http://www.intersurgical.co.uk/products/critical-care/connectors-and-elbows |
| F | Intersurgical Y-piece swivel connector 22M-22M-22M | http://www.intersurgical.co.uk/products/critical-care/connectors-and-elbows |
| G | Luer to hose barb adapter | e.g. https://www.coleparmer.co.uk/i/cole-parmer-male-luer-with-lock-ring-x-1-4-hose-barb-nylon-pack-of-25/4550519 |
| H | Hans Rudolf one-way valve | http://www.rudolphkc.com/img/uploads/pdf/691125%200714%20O.pdf |
| I | Intersurgical connector 22F-22F with inserted custom porous glass disc | e.g. https://www.heraeus.com/en/hca/products_hca/plates_quartz_1/filter_discs_1/filter_discs_hca.html |
| J | Intersurgical connector 22M-22M | http://www.intersurgical.co.uk/products/critical-care/connectors-and-elbows |
| K | Intersurgical tubing 22F 22mm | http://www.intersurgical.co.uk/products/anaesthesia/scavenging-tube |
| L | Intersurgical reservoir bag 2L 2F neck | http://www.intersurgical.co.uk/products/anaesthesia/aagbi-test-reservoir-bag |
| M | Fisher & Paykel humidifier | https://www.fphcare.co.nz/mr810/ |
| N | Intersurgical connector 22F-22F | http://www.intersurgical.co.uk/products/critical-care/connectors-and-elbows |
| O | Intersurgical Y-piece connector 22M-22M-22M | http://www.intersurgical.co.uk/products/critical-care/connectors-and-elbows |
| P | Intersurgical connector 22F-oxygen stem | http://www.intersurgical.co.uk/products/critical-care/connectors-and-elbows |
| Q | Gas Arc flowmeter 40 L/min | https://www.hisltd.co.uk/ProductDetail/Gas-Arc-Flowmeter/c67d8962-4ae6-4a97-8c7c-a9956d035ec8 |
| R | Bunzl healthcare 5mm bubble tubing | https://catalogue.bunzlhealthcare.co.uk/product/suction-bubble-tubing-non-sterile/ |
| S | Therapy Equipment railmounted carbon dioxide flowmeter | http://www.therapyequipment.co.uk/product/Diamond-Flowmeter-Range/products_flowmeters/index.html |
| T | Therapy Equipment railmounted oxygen flowmeter | http://www.therapyequipment.co.uk/product/Diamond-Flowmeter-Range/products_flowmeters/index.html |
| U | ADInstruments pressure transducer | https://www.adinstruments.com/products/bp-transducercable-kit |
| V | Intersurgical gas monitoring line | https://www.intersurgical.com/products/critical-care/gas-monitoring-lines |
| W | Buschjost Magnetventile GmbH & Co large bore solenoid valve for inspiratory loading | Type: Normally open 3/918-24/1002/R370-GN N.O. 24V DC<br>https://www.buschjostventile.de/attachments/1990a721-3afe-4099-a5e4-148763561294/de/Gewindeventile_112017.pdf<br>Plus required fittings (3 pcs.): MM052206N, John Guest Ltd. |
| X | Bürkert GmbH & Co solenoid valve for CO<sub>2</sub> | Type: Normally closed direct-acting 2/2-way valve, type 6013<br>https://www.burkert.co.nz/en/type/6013<br>Plus required fittings (one each): 1/8" BSP'P' (cone seat) swivel female x 1/4" BSPT' male adaptor, Part number FST-2BT-02-04, Custom Fittings Ltd., UK; British standard gas distinct direct fitting probe for 4 bar air with male 1/8" BSPP (cone seat) thread (part number 8502-01), Therapy Equipment, UK; British standard gas distinct single Schrader valve for 4 bar air with male 1/4" BSPT thread (part number 6102/P), Therapy Equipment, UK |
| Y | PowerBreathe device | https://www.powerbreathe.com/index.php |
| Z | Custom built controller box | See Figure 2 for details |
| ZA | TTL signal cables | DB25-to-BNC adapter (made in-house) and two generic BNC leads of appropriate length (e.g. part number L00015A1455, Telegärtner, for 10 metre leads) |
| ZB | Valve leads | 40881 Series cable assembly, part number 7000-40881-6360200, Murrelektronik |

### 2) Example Matlab code for the control of the circuit box

This code is suitable for use with a computer that has a parallel port, and requires the University of South Dakota Internet Psychology Lab’s MEX file plug-in (available from http://apps.usd.edu/coglab/psyc770/IO64.html). It is assumed that the TTL signals controlling the valves are taken from pins 2 and 3. If a computer without a parallel port is employed, a commercially available digital I/O-card may be used instead to produce the TTL signals, and the code adapted accordingly.

~~~
% Parallel port tester for valve control box
% Sebastian W Rieger
% Wellcome Centre for Integrative Neuroimaging (WIN)
% University of Oxford
% 02.05.2019
% Initialise port (if this fails, you need to install the mex-file plugin):
config_io;
% Set port address (you can find this in the Windows device manager):
port_address = hex2dec(‘378’);
% Energise the valve connected to pin 2 on the parallel port:
outp(port_address, 1);
% Wait for an appropriate amount of time (3 seconds in this example):
pause(3);
% De-energise the valves:
outp(port_address, 0);
% Energise the valve connected to pin 3 on the parallel port:
outp(port_address, 1);
% Wait for an appropriate amount of time (3 seconds in this example):
pause(3);
% De-energise the valves:
outp(port_address, 0);
~~~

